# Light-activated assembly of connexon nanopores in synthetic cells

**DOI:** 10.1101/2022.12.15.520663

**Authors:** Ahmed Sihorwala, Alexander Lin, Jeanne C. Stachowiak, Brian Belardi

**Affiliations:** McKetta Department of Chemical Engineering, University of Texas at Austin, Texas 78712; Department of Chemistry, University of Texas at Austin, Texas 78712; Department of Biomedical Engineering, University of Texas at Austin, Texas 78712

## Abstract

During developmental processes and wound healing, activation of living cells occurs with spatiotemporal precision and leads to rapid release of soluble molecular signals, allowing communication and coordination between neighbors. Non-living systems capable of similar responsive release hold great promise for information transfer in materials and site-specific drug delivery. One non-living system that offers a tunable platform for programming release is synthetic cells. Encased in a lipid bilayer structure, synthetic cells can be outfitted with molecular conduits that span the bilayer and lead to material exchange. While previous work expressing membrane pore proteins in synthetic cells demonstrated content exchange, user-defined control over release has remained elusive. In mammalian cells, connexon nanopore structures drive content release and have garnered significant interest since they can direct material exchange through intercellular contacts. Here, we focus on connexon nanopores and present activated release of material from synthetic cells in a light-sensitive fashion. To do this, we re-engineer connexon nanopores to assemble after post-translational processing by a protease. By encapsulating proteases in light-sensitive liposomes, we show that assembly of nanopores can be triggered by illumination, resulting in rapid release of molecules encapsulated within synthetic cells. Controlling connexin nanopore activity provides an opportunity for initiating communication with extracellular signals and for transferring molecular agents to the cytoplasm of living cells in a rapid, light-guided manner.

## Introduction

Responsive systems depend on information transfer to coordinate actions over long length scales. In living systems, molecular signals carry information from one cell to another, leading to collective, multicellular responses^1^ – a feat synthetic materials aim to emulate^2^. During paracrine signaling^3^, a major form of intercellular communication, a sender cell releases a diffusive molecular signal following stimulation to initiate signaling in surrounding receiver cells. To facilitate release of soluble signals, nanometer-diameter pores, carriers, and channels can act as conduits through the hydrophobic cell membrane to the extracellular space.^4^ The precise timing and extent of signal release is highly regulated to ensure that signaling in nearby cells occurs in a rapid manner without spurious activation or overactivation, which can lead to uncoordinated and deleterious effects.^5,6^ Engineering equivalent control over user-defined molecular release holds great promise for stimuli-responsive and spatially directed drug delivery^7^ and wound healing^8^ in vivo. Yet, significant obstacles remain as living cells employ a multitude of integrated mechanisms and pathways to keep content release under tight control, complicating engineering efforts.

Owing to their tunable composition and function, synthetic cells offer tremendous potential for responsive and controlled release.^9–12^ Both expression of multipass proteins in synthetic cell membranes^13–15^ and transport through the membrane^16–19^ have previously been demonstrated. However, without the regulatory machinery of cells, techniques for controlling synthetic cell responses must be designed and implemented. To activate release, synthetic cells require new innovations, such as mechanisms for triggering membrane pore activity, leading to controlled release. How to re-engineer membrane pores to enable rapid, stimuli-responsive release has thus garnered significant interest as of late, despite limited successes.^20–28^ Rather, recent work has focused on initiating synthetic cell leakage through other regulatory means.^29^ For instance, induced expression of a membrane pore, α-hemolysin,^30–34^ has seen success in turning ‘on’ synthetic cell leakage. While sophisticated, expression-dependent responses are often slow (hours timescale) due to the time-lag between transcriptional triggering and ultimate assembly of the translated pore protein.

Membrane pores come in various molecular conformations but can be grouped into two broad categories: single proteins^35^ and multimeric complexes^36^. Multimers assemble into a pore after translation and thus present an opportunity to engineer fast responses by post-translational processing. Connexins are one such family of membrane proteins. Connexins self-associate to form hexameric connexon nanopores that release cytoplasmic contents to the extracellular space and to neighboring cells^37,38^, offering an attractive option for synthetic information transfer and spatiotemporal control over cytoplasmic therapeutic delivery^39–43^. Fortunately, several pore structures of connexins have recently been solved by Cryo-EM, opening the door for structure-guided design of an activatable pore.^44–48^

Here we report the light-activated assembly of a connexon nanopore in synthetic cells and the subsequent release of internal contents. We re-engineer connexin-43 (Cx43)^46,47^ with a bulky protease recognition domain, which allows membrane-protein association but prevents nanopore function in the membrane. After enzymatic digestion, Cx43 rapidly assembles into a functional nanopore, releasing contents into the extracellular space. To provide user-defined control over the system, light-sensitive liposomes containing a protease are encapsulated in synthetic cells alongside the engineered Cx43. Light then acts as the stimulus to trigger Cx43 post-translational processing and nanopore assembly. Our data suggest that structure-guided engineering of membrane proteins is a fruitful strategy for incorporating responsiveness into synthetic cells, giving rise to controlled release.

## Results and Discussion

To explore engineering Cx43 nanopore activity, we set out to establish a synthetic cell system capable of Cx43 membrane incorporation and content release. Cx43-EGFP was cloned into a pRSET plasmid, and the plasmid along with machinery for transcription and translation (PURE system^49^) was encapsulated in vesicles using an inverted emulsion procedure^50^ (Fig. 1a). With this system, we first optimized Cx43 expression by varying the lipid composition (Fig. S1) and found that Cx43 expression was highest in POPC-containing vesicles compared to vesicles composed of either DOPC or DPPC. In POPC vesicles, we then monitored Cx43 expression over time and observed a plateau in the percentage of vesicles expressing Cx43 after a 2 h, 37 °C incubation (Fig. S2). Using POPC vesicles and a 2 h expression time as optimized parameters, we next imaged Cx43 localization in vesicles. Cx43 was solely localized to the membrane of POPC vesicles following expression (Fig. 1b), indicating that Cx43 associated with the lipid bilayer in the purified system. Membrane proteins can adopt either a non-functional interfacial conformation or a functional inserted conformation in the presence of lipid bilayers.^51,52^ Since Cx43 nanopores assemble from the inserted conformation, spontaneous insertion was tested by incubating vesicles with a peptide, Gap26, that recognizes one of the extracellular loops of Cx43.^53^ We found that Gap26 enriched on vesicle membranes with Cx43 expression but not on vesicle membranes lacking Cx43 (Fig. 1c), suggesting that Cx43 can adopt an inserted conformation after membrane association.

**Figure 1:**
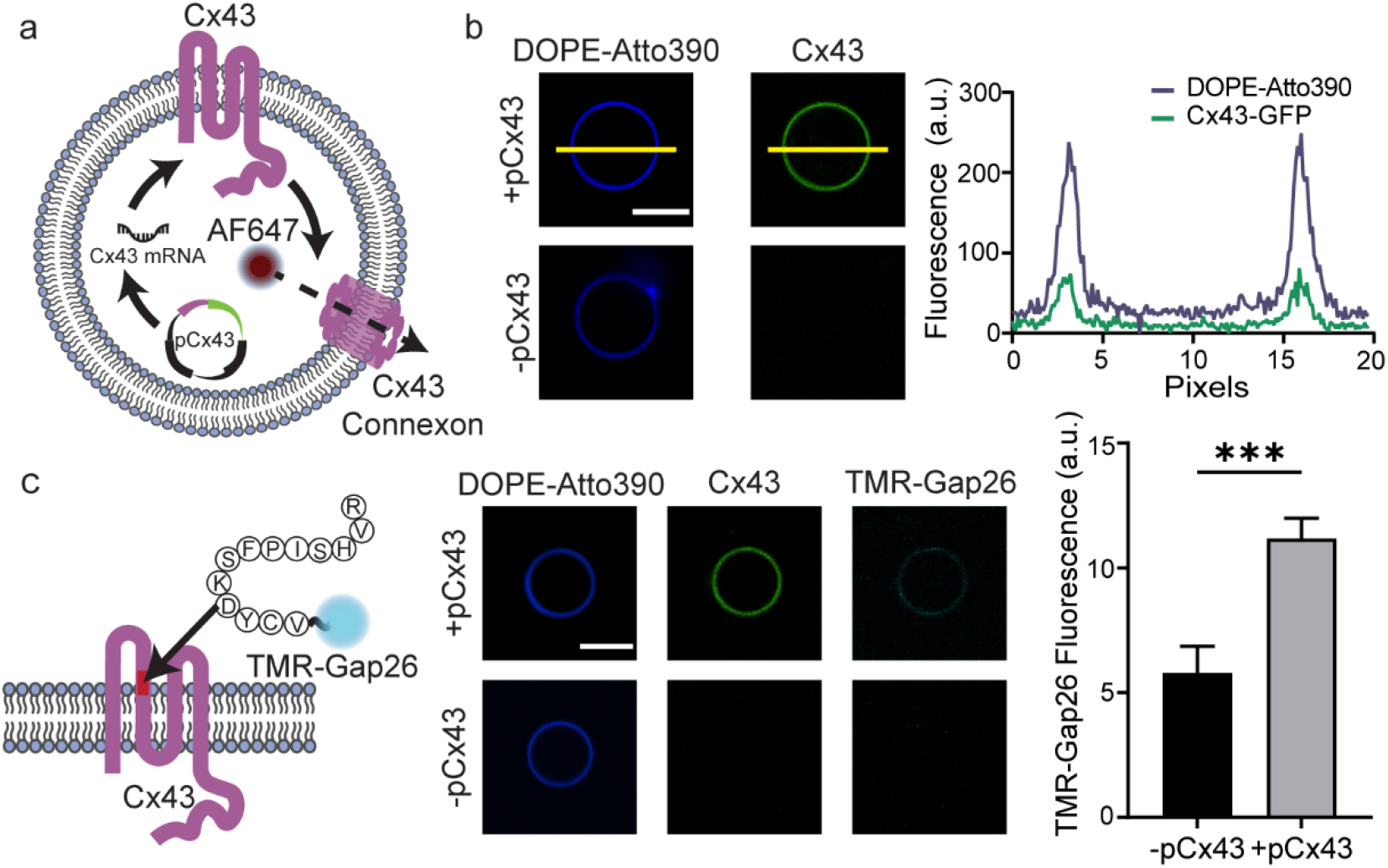
Membrane incorporation of Cx43 connexon nanopores. (a) Schematic of Cx43 connexon expression in synthetic cells. PURE system is co-encapsulated with Cx43-EGFP plasmid and AF647 dye in POPC/DOPE-Atto390 vesicles. Spontaneous membrane insertion of the expressed Cx43 protein is followed by the assembly of connexon nanopores that leads to AF647 dye leakage. (b) Fluorescence micrographs of vesicles encapsulating PURE system (left). Plot of DOPE-Atto390 and GFP fluorescence along yellow line indicates membrane association of Cx43 (right). (c) Schematic of TMR-Gap26 peptide binding the first extracellular loop of Cx43 (left, red). Fluorescence micrographs of vesicles incubated with TMR-Gap 26 peptide in the outer solution (center). Comparison of TMR membrane fluorescence between vesicles with and without Cx43 expression suggests membrane insertion of Cx43 (right). Error bars represent s.e.m. (*n* = 30 for -pCx43 and *n* = 45 for +pCx43). Scale bars: 10 µm.

In cells, connexon nanopores release ions, small molecules, and short polymers to the extracellular space.^54,55^ To examine whether Cx43 expressed in synthetic cells can perform the same function, we encapsulated a soluble small fluorescent molecule, AF647, in vesicles. After Cx43 expression, small molecule release was observed by fluorescence microscopy (Fig. 2). Release was rapid and correlated with the number of vesicles expressing Cx43 (Fig. S3), which agrees with a previous report for single α-hemolysin pores in 10 µm vesicles.^56^ Vesicles containing the PURE system but lacking Cx43 expression displayed a non-negligible amount of release, owing to defects in vesicles during the inverted emulsion procedure. In the future, background release might be addressed by alternative methods in vesicle production.^57–59^ Within living cells, Cx43’s pore activity is sensitive to the presence of extracellular calcium.^60^ We therefore asked whether Cx43 in purified membranes acted similarly to endogenous Cx43 pores by incubating vesicles with 2 mM calcium. We observed robust inhibition of pore activity in the presence of the divalent cation (Fig. S4). Within synthetic cells, Cx43 appeared to insert into pure membranes and assemble into a functional nanopore.

**Figure 2:**
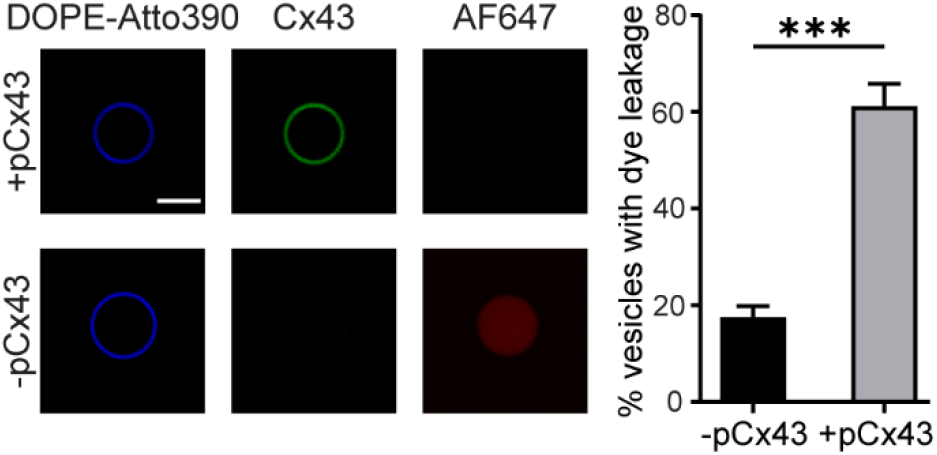
Content release from Cx43 connexon nanopores in synthetic cells. Fluorescence micrographs (left) and quantification of vesicles with AF647 dye leakage (right). Expression of connexon nanopores leads to vesicle leakage. Error bars represent the s.d. of 3 independent trials, at least 30 vesicles analyzed per trial. Scale bar: 10 µm. Asterisks represent statistically significant difference (two-tailed unpaired *t* test, *** *p <* 0.001).

With a functional Cx43 synthetic cell system, we turned our attention to suppressing Cx43 nanopore activity. Previous work in cells showed that Cx43 function is sensitive to cytoplasmic domain modification,^61^ and recent Cryo-EM structures of connexon pores,^44,45,48^ including Cx43’s,^46,47^ present an opportunity to use structure-guided design to engineer inhibition of Cx43. For Cx43, two solved structures show that the N-terminus provides key contacts at the bottom of the pore (cytoplasmic side) and faces inward toward the vacant channel (Fig. 3a). Since Cx43 assembles into a hexamer, Cx43’s collective N-termini exclude significant volume, leaving little free space. We reasoned that sterically encumbering the N-terminus would force Cx43 connexons to adopt a non-native conformation and consequently hinder pore activity. To test this hypothesis, we fashioned Cx43 with a bulky N-terminal group, the mCherry domain (Fig. 3b). After expression of the fused mCherry-Cx43 in vesicles, we observed a drastic reduction (69% decrease) in pore activity compared to wildtype Cx43 (Fig. 3c), indicating that bulky modifications to the N-terminus of Cx43 results in altered pore activity. Although impressive, further inhibition, and a larger dynamic range, might be possible by elucidating the effect of bulk dimensions on multimer assembly in membranes.^62–64^

**Figure 3.**
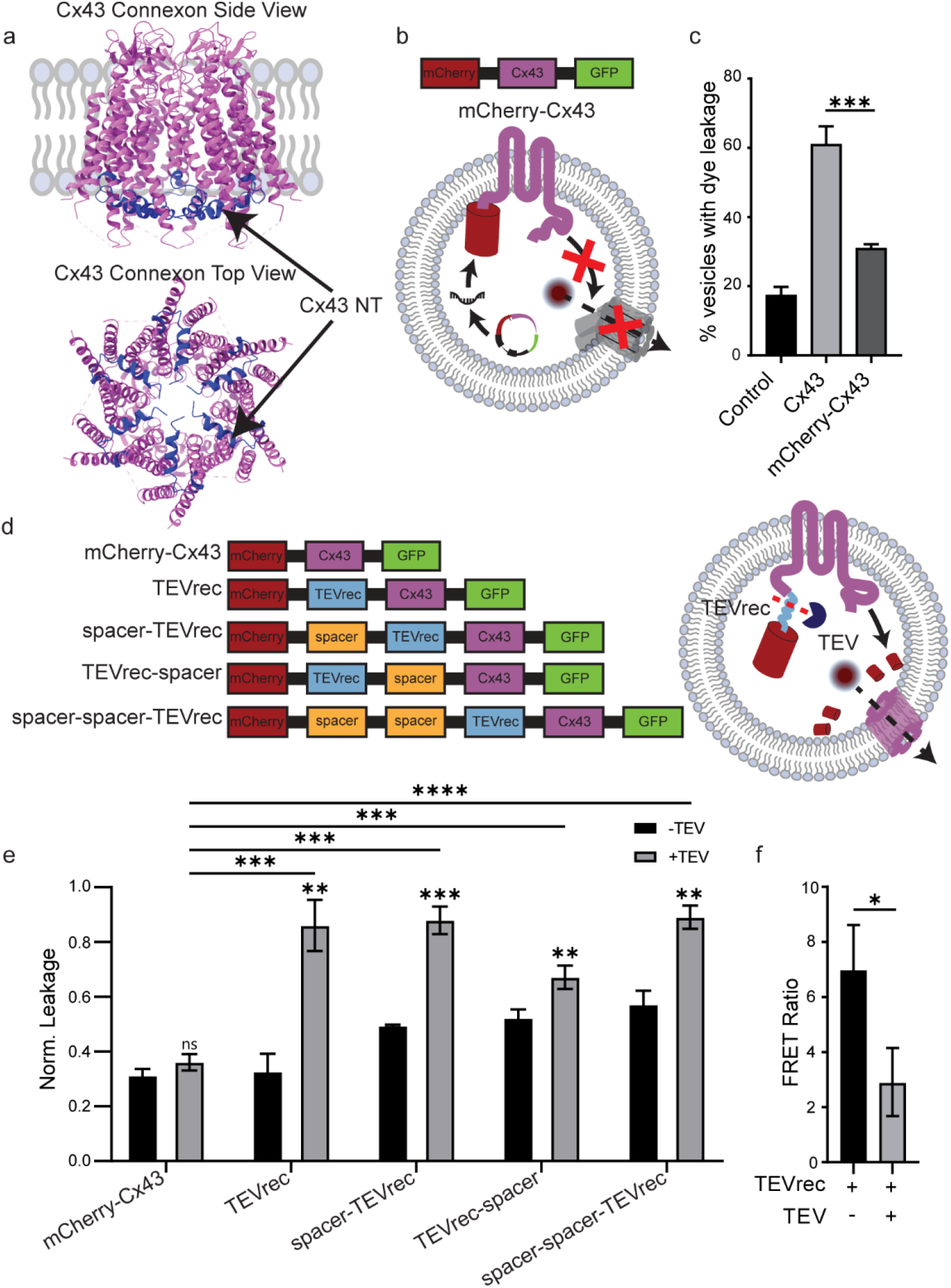
Controlling Cx43 connexon nanopore activity with protease-sensitive N-terminal bulk. (a) Side and Top views of Cx43 connexon, PDB: 7F93. The N-terminus of Cx43 monomers is highlighted in blue. (b) Schematic of mCherry-Cx43 expression and nanopore inactivity in synthetic cells. (c) Percentage of vesicles releasing dye after Cx43 or mCherry-Cx43 expression. N-terminal bulk limits pore activity. Error bars represent the s.d. of 3 independent trials, at least 30 vesicles were analyzed per trial. Asterisk represents statistically significant differences with Cx43 (two-tailed unpaired t test, *** p < 0.001). (d) Primary sequence of TEV protease-sensitive Cx43 variants (left). Schematic of connexon activity recovery with the addition of TEV protease (right). (e) Normalized dye leakage in vesicles expressing TEV-sensitive Cx43 variants in presence and absence of TEV. Error bars represent the s.d. of 3 independent trials, at least 30 vesicles were analyzed per trial. Asterisks represent statistically significant differences (two-tailed unpaired t test, ** p < 0.01, ***p < 0.001, **** p < 0.0001, n.s. p > 0.05). (f) FRET ratio of vesicles expressing TEVrec Cx43 variant in presence and absence of TEV. The lower FRET ratio for vesicles with TEV protease demonstrates successful mCherry cleavage. Error bars represent the s.d. of 3 independent trials. Asterisk represents statistically significant difference (two-tailed unpaired t test, * p < 0.05).

Encouraged by our results above, we next sought a strategy to dynamically remove the N-terminal constraint on Cx43 pore assembly, reversing the effects of steric exclusion and leading to assembly of functional pores, followed by content release. Protease processing would offer a rapid post-translation mechanism for removing the bulky group, and accordingly we examined whether TEV protease activity could transition Cx43 from an inactive to an active state. We designed a number of Cx43 variants containing an mCherry domain followed by the TEV recognition sequence (TEVrec) and either zero, one, or two peptide (GGGGS) spacers (Fig. 3d). Peptide spacers were included to provide flexibility in case TEV’s access to TEVrec was impeded by mCherry. We first expressed each variant in vesicles and tested dye leakage in the absence of TEV protease (Fig. 3e). A comparable and low extent of dye leakage was observed for both the TEVrec variant and mCherry-Cx43. The addition of spacer sequences resulted in modest increases in dye leakage, suggesting that spacers distance mCherry from the pore and limit its steric influence. We then repeated the experiment but included TEV protease in the lumen of vesicles. For each variant equipped with a TEVrec sequence, we found a significant increase in vesicle dye leakage compared to leakage in vesicles that lacked TEV (Fig. 3e). Cx43 without a spacer displayed the highest change in leakage (2.6-fold) in the presence of TEV and was used in subsequent experiments. To verify the TEVrec variant’s cleavage in vesicles, we measured Förster Resonance Energy Transfer (FRET). We found that FRET decreased in vesicles with the addition of TEV (Fig. 3f), which is expected since cleavage would increase the average distance between the acceptor (mCherry) and the donor (Cx43-EGFP). Together, these experiments demonstrate that protease cleavage can be used to assemble functional connexons from an inactive, sterically inhibited state in synthetic cells.

To regulate Cx43 pore activity, we lastly focused on rapid methods that could be used to ‘turn-on’ TEV processing. Light-mediated activation can act on fast time scales by inducing conformational changes^65–68^ and by initiating radical and carbene generation.^69–74^ Previously, the diyne lipid, DC(8,9)PC, has been shown to undergo photopolymerization, resulting in rupture of DC(8,9)PC-containing liposomes after minutes of UV light exposure.^75,76^ With this in mind, we decided to examine whether encapsulating TEV protease in photosensitive liposomes would lead to light-activated TEV processing. To test this idea, we first encapsulated TEV in 100 nm DC(8,9)PC-containing liposomes (Fig. S5) by hydrating a lipid film with a TEV solution and extruding the sample multiple times until a dense liposome suspension was formed. Next, we monitored TEV activity in the presence and absence of UV light. Without UV irradiation, DC(8,9)PC liposomes encapsulating TEV (DC(8,9)PC{TEV}) displayed minimal protease activity. In contrast, TEV activity from DC(8,9)PC{TEV} liposomes increased over time in the presence of 254 nm light with a t_1/2_ of ∼10 min (Fig. 4a), indicating liposome rupture and TEV processing occurs on the minutes timescale. We also verified that DC(8,9)PC{TEV} liposomes could be ruptured with light inside POPC vesicles (Fig. 4b). Specifically, we co-encapsulated DC(8,9)PC{TEV} liposomes and a TEV substrate labeled with an N-terminal fluorescent dye and C-terminal quencher inside POPC vesicles. Following UV irradiation, fluorescence dequenching of the TEV substrate (25.8-fold increase compared to non-irradiated sample) was observed, thereby confirming release of TEV from liposomes within vesicles.

**Figure 4.**
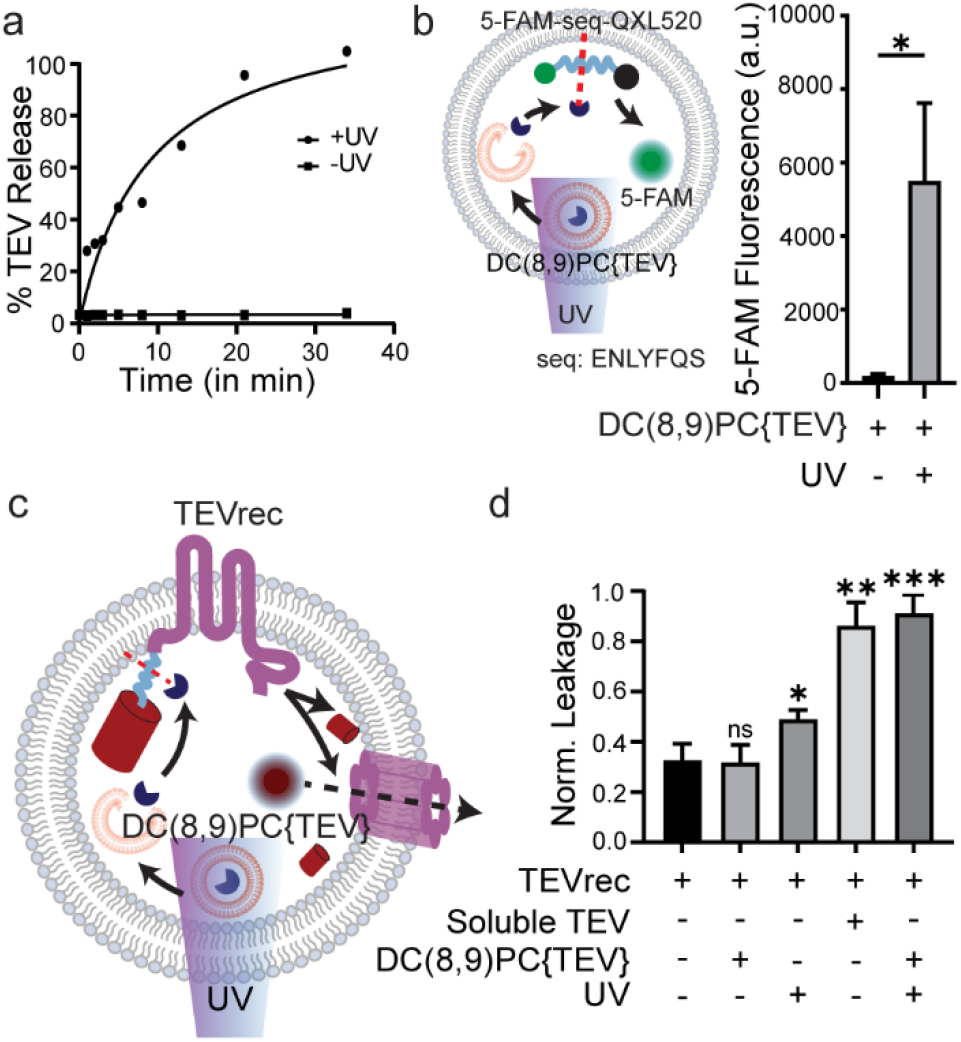
Light-activated assembly of functional connexon nanopores in synthetic cells. (a) Percentage of TEV release from DC(8,9)PC{TEV} liposomes as a function of illumination time (*n* = 2). (b) Schematic of light-mediated release of TEV from DC(8,9){TEV} liposomes encapsulated in synthetic cells (left). UV-responsive DC(8,9)PC{TEV} liposomes and a TEV substrate, seq, labeled with a fluorescent dye, 5-FAM, and a quencher, QXL520, were co-encapsulated inside POPC vesicles. Following illumination, photopolymerization of the liposomes leads to TEV release, resulting in the cleavage of the TEV substrate. Comparison of 5-FAM fluorescence in vesicles before and after 10 min UV irradiation (right). Error bars represent s.d. of 3 independent trials. Asterisk represents statistically significant difference (two-tailed unpaired *t* test, * p < 0.05). (c) Schematic of light-mediated assembly of functional connexon nanopores in synthetic cells. UV-responsive DC(8,9)PC{TEV} liposomes and AF647 were co-encapsulated in synthetic cells expressing TEVrec Cx43 variant. UV illumination results in TEV release from liposomes and cleavage of mCherry, leading to dye release. (d) Normalized dye leakage in vesicles expressing TEVrec Cx43 variant in the presence and absence of DC(8,9)PC{TEV} liposomes and UV illumination. Error bars represent the s.d. of 3 independent trials, at least 30 vesicles were analyzed per trial. Asterisks represent statistically significant differences with TEVrec only (two-tailed unpaired *t* test, *p < 0.05, ** p < 0.01, ***p < 0.001, n.s. p > 0.05).

With a method for triggering TEV processing, we tested the ability of light to initiate a rapid cascade that culminates in Cx43 nanopore assembly and synthetic cell release. To do this, we produced vesicles expressing the TEVrec variant of Cx43 and containing DC(8,9)PC{TEV} liposomes (Fig. 4c). In the absence of light, Cx43 pore activity remained at background levels (Fig. 4d) as expected for an inactive Cx43 state. Gratifyingly, after exposing vesicles to 10 min (t_1/2_) of 254 nm light, we observed a 2.9-fold increase in Cx43 pore activity. In fact, the amount of leakage after UV exposure was indistinguishable from TEVrec-expressing vesicles with soluble TEV, suggesting that light-activated release of TEV was efficient. Our results suggest that light-activation of Cx43 assembly can serve as a rapid timescale responsive system and lead to content release across bilayer structures.

## Conclusion

In sum, we have developed a rapid method for controlling the functional assembly of a nanopore in synthetic cells. To accomplish this goal, we rationally designed a bulky modification that sterically disrupts the assembly of functional Cx43 nanopores. By engineering the modification as a cleavable substrate, enzymatic processing was harnessed to transition the pore from an ‘off’ state to an ‘on’ state. As most methods display significant time lags between activation and pore activity,^30–34,77^ we sought to build a rapid responsive system capable of content exchange. By linking enzymatic activity to light exposure, our synthetic cell system was responsive on fast timescales and exhibited subsequent molecular information exchange. In the future, this strategy may prove useful for engineering other membrane pores to be responsive, since the termini of many pore proteins contribute key contacts for functionality.^78–80^ As well, this form of responsive pore formation can be implemented to control the precise time and location of information transfer with applications in drug delivery and engineering coordinated activity in both biotic and abiotic environments.

## Acknowledgement

This work was supported by grants from the National Institutes of Health (R35GM142941 to B.B. and R35GM139531 to J.C.S.), the National Science Foundation (Grant 2218467 to B.B. and J.C.S.), and the Welch Foundation (Grant F-2055-20210327 to B.B. and Grant F-2047 to J.C.S.). A.L. is supported by an NSF predoctoral fellowship.

