## Supporting Information for "Light-activated assembly of connexon nanopores in synthetic cells"

*Affiliations*

### Materials and Methods

General Methods

All of the chemical reagents were of analytical grade, obtained from commercial suppliers and used without further purification, unless otherwise noted. 4-(2-hydroxyethyl)-1-piperazineethanesulfonic acid (HEPES), sodium chloride (NaCl), calcium chloride ($CaCl_2$) and OptiPrep Density Gradient Medium were purchased from Sigma-Aldrich.1-palmitoyl-2-oleoyl-glycero-3-phosphocholine (POPC), 1,2-dioleoyl-*sn*-glycero-3-phosphocholine (DOPC), 1,2-dipalmitoyl-*sn*-glycero-3-phosphocholine (DPPC), 1,2-bis(10,12-tricosadiynoyl)-*sn*-glycero-3-phosphocholine (DC(8,9)PC), 1,2-distearoyl-*sn*-glycero-3-phosphoethanolamine-N-[methoxy(polyethylene glycol)-2000] (DSPE-PEG(2000)) and Mini Extruder kit were obtained from Avanti Polar Lipids. 1,2-Dioleoyl-*sn*-glycero-3-phosphoethanolamine labeled with Atto 390 (DOPE-Atto390) was obtained from ATTO-TEC GmbH. Tetramethylrhodamine{Ahx}VCYDKSFPISHVR (TMR-Gap26) was purchased from GenScript Biotech. Alexa Fluor 647-NHS Ester was obtained from ThermoFisher Scientific. TEV Protease was purchased from New England Biolabs. Sensolyte 520 TEV Protease Assay Kit Fluorimetric was obtained from AnaSpec.

Fluorescence imaging was carried out on an Eclipse Ti2 microscope (Nikon) equipped with 405/488/560/640nm lasers (Andor), a Dragon 500 high speed spinning disk confocal module (Andor), and a Zyla 4.2 sCMOS camera (Andor). Fluorescence micrographs of vesicles were acquired with a 60x objective (Nikon, NA 1.49 TIRF) and were analyzed using Fiji (ImageJ). Brightness/contrast for all analyzed channels was kept consistent across images within a single experiment. Sample fluorescence was acquired with a BioTek Cytation 5 Imaging reader. For UV irradiation experiments, samples were placed in a 96-well plate at a distance of 6.5 cm from a UVP 3UV lamp (254 nm shortwave UV emission, 115V, 8W, Analytikjena).

Plasmid constructs

Cloning was performed using routine procedures. The Cx43-EGFP fusion gene was obtained by amplifying the Cx43[1] and EGFP genes using PCR and introducing them into the pRSET vector between XbaI and EcoRI with Gibson Assembly (plasmid pRSET_Cx43-EGFP). The Cx43 variants were designed in a similar way by amplifying the Cx43-EGFP fusion gene and mCherry gene using PCR and inserting them between the XbaI and EcoRI sites of the pRSET vector using Gibson Assembly (pRSET_mCherry-Cx43-EGFP, pRSET_mCherry-TEVrec-Cx43-EGFP, pRSET_mCherry-spacer-TEVrec-Cx43-EGFP, pRSET_mCherry-TEVrec-spacer-Cx43-EGFP, pRSET_mCherry-spacer-spacer-TEVrec-Cx43-EGFP). The oligonucleotides used to introduce the spacers and the TEV recognition site for the TEVrec, spacer-TEVrec, TEVrec-spacer and spacer-spacer-TEVrec variants are listed in the following table (highlighted bases represent hybridization sites for PCR):

| Cx43 variant | Oligonucleotide sequence (5’-3’) |
|---|---|
| TEVrec | Forward:<br>AGAAAACCTGTATTTTCAGAGCATGGGTGACTGGAGTGCC<br>Reverse:<br>TCACCCATGCTCTGAAAATACAGGTTTTCTTTGTATAATTCGTCCATTCCACCTG |

| | |
|---|---|
| spacer-TEVrec | Forward: GCGGGGGCAGCGAAAACCTGTATTTTCAGAGCATGGGTGACTGGAGTGCC<br>Reverse: TCTGAAAATACAGGTTTTCGCTGCCCCCGCCACCTTTGTATAATTCGTCCATTCCACCTG |
| TEVrec-spacer | Forward: ACCTGTATTTTCAGAGCGGTGGCGGGGGCAGCATGGGTGACTGGAGTGCC<br>Reverse: TGCCCCCGCCACCGCTCTGAAAATACAGGTTTTCTTTGTATAATTCGTCCATTCCACCTG |
| spacer-spacer-TEVrec | Forward: GGCAGCGGTGGAGGAGGTAGTGAAAACCTGTATTTTCAGAGCATGGGTGACTGGAGTGCC<br>Reverse: CAGGTTTTCACTACCTCCTCCACCGCTGCCCCCGCCACCTTTGTATAATTCGTCCATTCCACCTG |

Vesicle synthesis and Cx43-GFP expression

Vesicles with Cx43-GFP expression were prepared using a modified inverted emulsion process.[2] A 657 µM (total lipid) suspension of POPC:DOPE-Atto390 (99.9:0.1) in a silicone oil, mineral oil and decane (40:7:3) mixture was freshly prepared by sonication (30 min) before each experiment. 20 µL of this lipid-oil solution was added on top of a 90 µL 5 mM HEPES and 30 mM NaCl solution (pH 7.4) in a 1.5 mL tube and allowed to rest for 30 min. 7 µL of the inner vesicle solution containing PURExpress system (solutions A and B), Cx43-EGFP plasmid (pCx43, 70 ng), and 10% OptiPrep was prepared at 4 °C in buffer. 5 µL of the inner solution was added to 80 µL of the lipid-oil solution and vortexed for 50 s to create an emulsion. Next, the emulsion was gently layered over the lipid monolayer interface in a 1.5 mL tube. Centrifugation was immediately performed at 4 °C for 10 min at 2400 x *g*. After centrifugation, the tube was punctured at the bottom, and the vesicles in buffer were ejected into a fresh 1.5 mL tube. This tube was incubated at 37 °C for 2 hours to facilitate Cx43-EGFP expression. Subsequently, the prepared vesicles expressing Cx43-EGFP were imaged using a 60x objective. A similar protocol was used for all subsequent experiments.

To study the effect of lipid on Cx43 expression, the above procedure was repeated in vesicles with the following membrane compositions: DOPC:DOPE-Atto390 (99.9:0.1) and DPPC:DOPE-Atto390 (99.9:0.1). The vesicles were then imaged using a 60x objective and the acquired fluorescence micrographs were analyzed in Fiji (ImageJ). Briefly, multiple line scans were performed to obtain the average GFP fluorescence at the membrane for each vesicle while keeping the contrast/brightness for the GFP channel consistent across all conditions.

A time course for Cx43 expression in vesicles was conducted by incubating vesicles at 37 °C for different durations (10, 30, 60, 120 and 180 min respectively) and subsequently imaging these vesicles using a 60x objective at different time points. Percentage of vesicles with Cx43 expression at each time point was calculated by observing membrane GFP fluorescence using Fiji (ImageJ).

Gap26 peptide assay

To test membrane insertion of Cx43, vesicles were prepared using the above vesicle formation method with the exception that the outer solution also contained 1 μM of TMR-Gap26 peptide in buffer (5 mM HEPES and 30 mM NaCl, pH 7.4). The control vesicles lacked the Cx43 plasmid (-pCx43) in the inner solution. The prepared vesicles were imaged using a 60x objective with identical settings for ±pCx43. The acquired fluorescence micrographs were analyzed in Fiji (ImageJ). Briefly, multiple line scans were performed to obtain the average TMR fluorescence at the membrane for each vesicle while keeping the contrast/brightness for the TMR-Gap26 channel consistent across ±pCx43 conditions.

Vesicle leakage assay

To test the assembly of functional Cx43 connexons in vesicles, 10 μM Alexa Fluor 647-COOH was added to the inner solution mix. Control vesicles encapsulating AF647 but lacking the Cx43 plasmid (-pCx43) in the inner solution were also prepared in a similar way. The prepared vesicles were imaged using a 60x objective with the same settings for ±pCx43. The acquired fluorescence micrographs were analyzed in Fiji (ImageJ) to quantify percentage of vesicles with dye leakage. Vesicles with lumenal dye fluorescence less than 10% relative to background fluorescence were determined using the oval selection tool in Fiji and classified as vesicles with dye leakage.

A time course of dye leakage from Cx43 expressing vesicles was performed by incubating vesicles at 37 °C for different durations (10, 30, 60, 120 and 180 min respectively) and imaging vesicles at each time point using a 60x objective. Subsequently, percentage of vesicles with dye leakage at each time point was calculated.

To investigate $Ca^{2+}$ sensitivity of the assembled Cx43 connexons in vesicles, 2 mM $CaCl_2$ in buffer (5 mM HEPES and 30 mM NaCl, pH 7.4) constituted the outer vesicle solution. Vesicles were imaged using a 60x objective, and percentage of vesicles with dye leakage was quantified and compared to Cx43 expressing vesicles that lacked $CaCl_2$ in the outer solution.

For connexon inhibition experiments, the above dye leakage analysis was performed by expressing the various Cx43 variants (mCherry-Cx43, TEVrec, spacer-TEVrec, TEVrec-spacer, spacer-spacer-TEVrec) in vesicles using the PURExpress system and their respective plasmids. Again, vesicles were imaged using a 60x objective with the same settings for each Cx43 variant and subsequently analyzed using Fiji (ImageJ) to quantify percentage of vesicles with dye leakage. These values were then normalized with respect to the -pCx43 (0) and +pCx43 (1) controls to calculate fold-inhibition.

To test TEV protease sensitivity of the designed Cx43 variants, vesicles expressing a single Cx43 variant and encapsulating TEV protease (5 Units) were subject to the above dye leakage assay. Following image acquisition and analysis, percentage of vesicles with dye leakage were normalized with respect to -pCx43 (0) and +pCx43 (1) controls to calculate fold-recovery.

FRET Assay

Vesicles expressing the Cx43 TEVrec variant were prepared with and without TEV protease in the inner solution according to the vesicle synthesis procedure above. 20 µL of vesicle solution was added to a 384 well plate (Greiner Microplate 384-well, Black) after a 2 hour, 37 °C incubation period. Vesicles expressing Cx43 and vesicles lacking the Cx43 plasmid served as controls. End-point fluorescence readings for mCherry ($\lambda_{ex/em}$ = 587/610 nm), GFP ($\lambda_{ex/em}$ = 488/507 nm) and FRET channels ($\lambda_{ex/em}$ = 488/610 nm) were obtained using the plate reader. No plasmid fluorescence values were subtracted from the TEVrec conditions and Cx43 control. The TEVrec conditions (±TEV) were then normalized with respect to the Cx43 fluorescence values to account for GFP bleed-through. Finally, the ratio of the FRET channel fluorescence to GFP fluorescence for TEVrec was calculated.

Preparation of DC(8,9)PC{TEV} liposomes

A well-mixed solution of DPPC:DC(8,9)PC:DSPE-PEG2000 (79.5:20:0.5) lipids in chloroform (10 mg/mL) was placed under a stream of $N_2$ gas to evaporate the chloroform. The film was kept in a vacuum desiccator for 2 hours to rid the lipid cake of any residual chloroform. Films containing DC(8,9)PC were prepared in the absence of light and covered with foil. Next, the lipid film was hydrated with an aqueous solution containing 100 Units of TEV protease, 1 mM DTT, 25 mM HEPES, 150 mM NaCl, pH 7.4, at 42 °C to form multilamellar vesicles. Multilamellar vesicles were then extruded 43 times through a 100 nm polycarbonate filter at 42 °C to prepare liposomes encapsulating TEV protease, and the resulting liposomes were kept at 4 °C for 30 minutes. Finally, liposome samples were centrifuged at 16900 x *g* for 10 min at room temperature to separate any large lipid aggregates from liposomes.[3]

Dynamic Light Scattering of liposomes

Liposome diameter was determined via dynamic light scattering using a Zetasizer Nano ZS size analyzer (Malvern Panalytical). 50 µL of liposome solution was diluted in 950 µL of 25 mM HEPES, 150 mM NaCl, pH 7.4 buffer, and size distribution of particles was measured via three runs and 16 measurements per run.

Kinetics of TEV leakage from prepared liposomes

The prepared DC(8,9)PC{TEV} liposomes were diluted 7-fold in a 25 mM HEPES, 150 mM NaCl, pH 7.4 buffer. Controls with unencapsulated soluble TEV were also prepared. 100 µL of DC(8,9)PC{TEV} liposome samples and controls were added to a 96-well plate (Corning 96-well, Black) and illuminated with 254 nm UV light for different durations (1, 2, 3, 5, 8, 13, 21 and 34 min). A non-UV illuminated aliquot of DC(8,9)PC{TEV} liposomes was kept at room temperature to determine TEV encapsulation efficiency in prepared liposomes. 0.5 µL of the TEV substrate (5-FAM-ENLYFQS-QXL 520 quencher, 100X) was diluted to 50 µL in buffer (25 mM HEPES, 150 mM NaCl, pH 7.4) and then added to each of the UV and non-UV irradiated samples, and the mixtures were allowed to incubate at 30 °C for 1 hour. Following the incubation step, end point 5-FAM fluorescence values were obtained using a plate reader ($\lambda_{ex/em}$

= 492/518 nm). The percentages of TEV release for the UV and non-UV irradiated DC(8,9)PC{TEV} liposomes were calculated by comparing their fluorescence values to that of their respective controls.

TEV release in vesicles

DC(8,9)PC{TEV} liposomes along with 25 mM HEPES, 150 mM NaCl, pH 7.4, 10% OptiPrep and 0.7 μL of the TEV substrate (10X) formed 7 μL of the inner solution of vesicles prepared using the above inverted emulsion procedure. Controls were prepared with no DC(8,9)PC{TEV} liposomes in the inner solution . Vesicles encapsulating DC(8,9)PC{TEV} liposomes and controls were irradiated with 254 nm UV light for 10 min followed by incubation at 30 °C for 1 hour. Simultaneously, aliquots of non-UV irradiated vesicles with and without liposomes were also incubated at 30 °C for 1 hour. Subsequently, end point 5-FAM fluorescence values for all samples and controls were analyzed with a plate reader ($\lambda_{ex/em}$ = 492/518 nm). The fluorescence values of the UV and non-UV irradiated samples were compared following the subtraction of their respective control fluorescence.

Light-activated assembly of connexon nanopores

DC(8,9)PC{TEV} liposomes along with the PURExpress system, TEVrec plasmid (70 ng), 10% OptiPrep and 10 μM AF647-COOH in buffer (5 mM HEPES, 30 mM NaCl, pH 7.4) were mixed to form 7 μL of the inner solution . These vesicles were synthesized with the inverted emulsion procedure above and incubated at 37 °C for 2 hours. The vesicles were then illuminated with 254 nm UV light for 10 min. Vesicles prepared similarly but not exposed to UV light or not encapsulating DC(8,9)PC{TEV} liposomes served as controls. UV-illuminated vesicles with DC(8,9)PC{TEV} liposomes and TEVrec expression were imaged using a 60x objective and subsequently analyzed for AF647 dye leakage using Fiji (ImageJ). The TEV and UV controls were also analyzed similarly.

### Supporting Information Figures

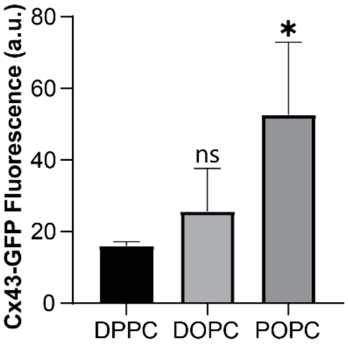


**Figure S1.** Cx43 expression in vesicles with different lipid compositions. Quantification of GFP membrane fluorescence in vesicles expressing Cx43. Highest Cx43 expression was found for the POPC membrane. Error bars represent the s.d. of 3 independent trials, at least 20 vesicles were analyzed per trial. Asterisks represent statistically significant differences with DPPC (two-tailed unpaired *t* test, *$p < 0.05$, n.s. $p > 0.05$).

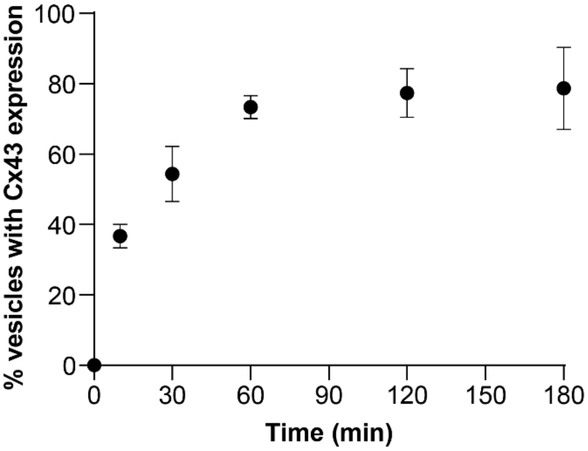


**Figure S2.** Time course of Cx43 expression in synthetic cells. Plot of percentage of vesicles with Cx43 expression as a function of incubation time at 37 °C. A plateau in Cx43 expression is observed after 2 h of incubation time. Error bars represent the s.d. of 3 independent trials, at least 20 synthetic cells were analyzed at each time point per trial.

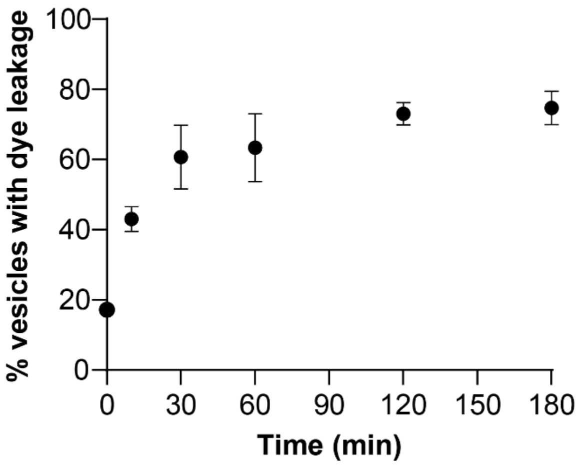


**Figure S3.** Time course of AF647 dye release in synthetic cells expressing Cx43. Plot of percentage of dye leakage in Cx43-expressing vesicles as a function of time at 37 °C. A plateau in percentage of dye leakage is observed after 2 h of incubation time, similar to Cx43 expression. Error bars represent the s.d. of 3 independent trials, at least 20 synthetic cells were analyzed at each time point per trial.

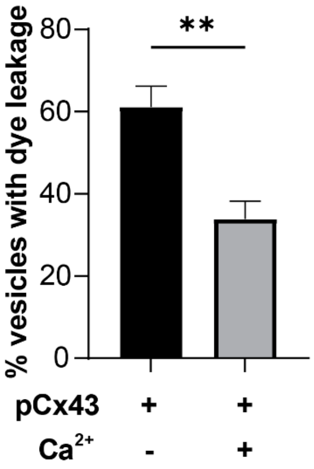


**Figure S4.** Effect of calcium on connexon nanopore activity. Percentage of synthetic cells with Cx43 expression releasing dye with and without 2 mM $Ca^{2+}$. Error bars represent the s.d. of 3 independent trials, at least 25 vesicles were analyzed per trial. Asterisks represent statistically significant differences (two-tailed unpaired *t* test, ** $p < 0.01$).

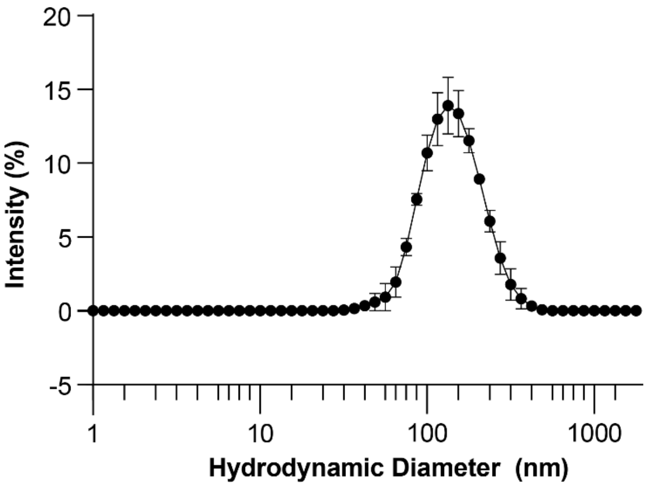


**Figure S5.** Characterization of DC(8,9)PC{TEV} liposomes by Dynamic Light Scattering. Error bars represent s.e.m. (*n* = 3).
